# Longitudinal Investigation of Structural and Resting-State Effective Connectivity Alterations in a Non-Human Primate Model of Huntington’s Disease

**DOI:** 10.1101/2025.07.18.665554

**Authors:** Gabriele Petraityte, Joëlle van Rijswijk, William A Liguore, Jodi L McBride, Marleen Verhoye, Daniele Bertoglio, Alison R Weiss, Mohit H Adhikari

## Abstract

Huntington’s disease (HD) is a genetic neurodegenerative disorder caused by expanded CAG repeats in the huntingtin gene which produce a mutant huntingtin (mHTT) protein that contributes to progressive striatal, cortical, and white-matter atrophy, resulting in motor dysfunction and cognitive decline. Recently, a non-human primate (NHP) model of HD was developed via stereotaxic delivery of an adeno-associated viral vector expressing 85 CAG repeats (85Q) into the striatum. This model recapitulates several neuropathological changes and symptoms observed in people with HD (PwHD) including chorea and mild cognitive impairment. A previous longitudinal, multimodal MRI investigation in this model revealed volumetric and resting-state functional connectivity (rs-FC) changes compared to controls, in key regions involved in HD, over the course of 30 months.

We aimed to study longitudinal changes in structural connectivity (SC), obtained from diffusion MRI scans from the same animals, comparing the 85Q animals to the control (Buffer) group. Additionally, going beyond the correlative rs-FC analyses, we investigated changes in causal, inter-regional functional interactions by estimating effective connectivity (EC) from rs functional MRI scans, constrained to strong structural connections. We found that the SC between basal ganglia regions and the cortex was reduced in the 85Q primates compared to the Buffer group at 14-months post virus injection, aligning with the pathological process observed in PwHD at later stages of the disease. EC from the caudate and putamen to the motor cortex was significantly reduced in the 85Q animals as early as 3-months post-injection providing novel insights into early alterations in causal functional interactions.

## 1. Introduction

Huntington’s Disease (HD) is a progressive autosomal dominant neurodegenerative disease caused by an expansion of CAG sequence (>40 repeats) in the huntingtin (*HTT*) gene. Repeat expansion codes for an elongated polyglutamine tract in the huntingtin protein (mHTT) affecting the neurons, in particular, GABAergic medium spiny neurons [1] of the striatum, resulting in neurotransmission changes [2] within the basal ganglia circuit. This leads to early symptoms such as chorea, impaired coordination and voluntary movement control and, as the disease progresses, movement rigidity and cognitive decline [3].

While a cure for the disease remains elusive, research aimed at advancing therapy necessitates the utilization of animal models for development of biomarkers capable of assessing and monitoring the efficacy of emerging therapeutic strategies. Although the mouse is one of the most used species in HD research, larger species such as non-human primates (NHP) help fill a translational gap by more accurately recapitulating key features of the disease. NHPs exhibit complex fine motor behaviors that closely mirror those observed in humans, allowing for more sensitive assessments of motor impairments. Additionally, their larger and more human-like brain anatomy—especially in regions critically affected by HD such as the striatum, motor and prefrontal cortices—offers a significant advantage. Unlike rodents, NHPs possess an internal capsule, a key white matter structure relevant to HD pathology, and a more developed prefrontal cortex, supporting higher-order cognitive functions. Their longer lifespan compared to rodents also enables longitudinal studies that can capture prodromal and early disease stages more in depth [4–6]. To address these translational challenges and better model the complexity of HD in humans, a novel rhesus macaque model was designed to mimic key aspects of the disease. This model carries a fragment of the human HTT gene with 85 CAG repeats (85Q), via injection of an adeno-associated viral vector into the caudate and putamen, expressed throughout cortico-basal ganglia circuit [6].

A multimodal MRI and behavioral investigation in this model, along with a Buffer group, which received a buffered saline injection, and a 10Q group, which expressed 10 CAG repeats, carried out longitudinally at the baseline (prior to virus injection surgery), and at 3-, 6-, 9-, 14-, 20-, and 30-months post-surgery, showed several changes in the 85Q animals. Additionally, two PET studies detected mHTT aggregates as well as alterations in dopaminergic D2/3 receptors and glucose metabolism in several brain regions in 85Q animals compared to the Buffer group [7, 8]. 85Q animals recapitulated the cognitive and progressive motor decline, including chorea, observed in PwHD [6]. Anatomical MRI revealed mild tissue atrophy in caudate, putamen, globus pallidus, frontal and motor cortex of the 85Q group compared to the Buffer group. Diffusion tensor imaging (DTI) MRI showed reduced fractional anisotropy (FA) in caudate, prefrontal, motor, temporal and parietal cortices as well as increased FA in the areas of putamen and globus pallidus as early as 3-months post-surgery persisting throughout the 30-month period [9] in the 85Q group. The independent component analysis (ICA) of resting-state functional MRI (rs-fMRI) data indicated reduced correlation between sensory-motor areas and the component comprising of the caudate, putamen, ventral sensory motor cortex, and medial prefrontal cortex in the 85Q group at multiple timepoints post-surgery [6]. In comparison, while it did not significantly differ from the Buffer group in most MRI and PET outcomes, the 10Q group did exhibit symptoms such as spatial working memory deficits, impaired fine motor skill performance, abnormal forelimb posture, forelimb tremor, and orofacial chorea at different timepoints [10].

These findings demonstrate regional volumetric, microstructural as well as inter-regional correlative functional alterations exhibited by the 85Q model animals. However, they do not reveal disconnections of inter-regional structural connectivity (SC) or changes in causal, directed functional interactions. Structural connectivity (SC) is a measure of strength of white matter tracts between pairs of grey matter regions, typically measured using diffusion MRI and tractography in humans, and shows strong relationship with rs-FC [11, 12]. It is also instrumental in estimating effective connectivity (EC) that uses SC as a prior to capture causal, directed functional interactions between brain regions from their rsfMRI BOLD signals. The compelling advantage of the EC approach lies in its capability to interpret changes in BOLD signals by incorporating anatomically plausible connections and distinguishes it from FC that only yields correlative information [13]. Recent studies have shown that EC can not only provide mechanistic explanations for emergence of FC but can be a more sensitive measure than FC as a fingerprint of individuals as well in classifying patients of neurological disorders [13–16]. To our knowledge, no study has investigated EC changes in animal models of HD, or in PwHD.

Therefore, in this study, we aimed to first investigate longitudinal SC changes in the 85Q model. Then, by using strongest structural connections in each individual animal as an anatomical prior, we inferred the EC from the individual’s rsfMRI data. We hypothesized that the expression of the fragment carrying 85 CAG repeats in the first three exons of the 85Q model animals leads to alterations in SC in brain regions affected by HD and/or in the regions of viral vector distribution. Moreover, since EC has been shown to be a more sensitive marker than FC in other neurological disorders [16], we hypothesized that early changes in causal functional interactions in this model will be captured by EC.

## 2. Materials and Methods

### 2.1 Animal Model

The data utilized in this study were acquired by Weiss et al. [6]. A total of 17 (n = 12 female, n = 5 male) adult (age 6-13 years) *Rhesus Macaques (Macaca mulatta)* were enrolled in the study. The animals were divided into 3 groups, the first group (n = 6) received a 1:1 mixture of recombinant adeno-associated viral vectors AAV2 and AAV2.retro expressing a glutamine encoded repeat with 85 pure CAG repeats, followed by a single CAA/CAG cassette (85Q). A second group (n = 6) was injected with equivalent mixture of viral vectors expressing a glutamine encoded repeat with 10 pure CAG repeats, followed by a single CAA/CAG cassette (10Q). The third group – Buffer, received buffered saline injection. Injections were made into the caudate (90μL pre-commissural and 60μL post-commissural) and putamen (95μL pre-commissural and 85μL post-commissural), with a per-hemisphere volume totaling 330μL. The injection sites were positioned at intervals of approximately 4-5mm from each other within both the caudate and putamen regions [6].

The 10Q group did express the 10 CAG repeats fragment but did not display formation of aggregates [8]. Compared to the Buffer group, it had a partly higher glucose uptake possibly reflecting neuronal susceptibility imbalance due to the excess 10Q protein presence [7]. Moreover, it differed significantly in working memory performance and motor deficits with both the 85Q as well as the Buffer group [6]. Therefore, we focused this study on investigating longitudinal SC and EC changes in the 85Q group compared to the Buffer group. Nevertheless, for completeness, we also carried out statistical comparisons between all three groups as described in the supplementary materials.

### 2.2 MRI acquisition

MRI scans were acquired prior to the surgery and at 3-, 6-, 9-, 14-, 20-, and 30-months after the surgery. For the duration of the scanning, all animals were anesthetized (ketamine HCl (15mg/kg intramuscular) for induction and 1-2% isoflurane gas vaporized in 100% oxygen for anaesthesia maintenance). The imaging procedures were conducted utilizing a Siemens Prisma whole body 3T MRI system with a 16-channel pediatric head RF coil. Four MRI modalities were obtained: T1-weighted MRI, T2-weighted MRI, diffusion-tensor imaging (DTI), and rs-fMRI with the total scan time of 122min 10s [6]. For this study only DTI and rs-fMRI scans were used. Diffusion tensor MRI data were obtained using a Diffusion-weighted (DW) spin-echo EPI sequence with phase-encoding along the anterior-to-posterior direction and isotropic voxels of (1.0 mm) and TR/TE=6700/73ms (GRAPPA factor = 2). Diffusion-weighted imaging (DWI) volumes with single b = 1000*s*/*mm*^2^ in 30 diffusion directions and seven repetitions of 6 b = 0 s/mm^2^ volumes were acquired. Additionally, a single b = 0 s/mm^2^ volume was acquired in the reverse phase-encoding direction to later correct for susceptibility-induced distortions. For rs-fMRI, 784 3D volumes of BOLD images were acquired with T2*-weighted gradient echo-EPI sequence with TE/TR = 25/2290 ms, flip angle = 79°, and isotropic voxels of 1.5 mm^3^ [6].

### 2.3 Parcels Assignments

For the SC and EC estimation, 55 grey-matter ROIs from both hemispheres spanning the whole brain were used, as defined by the ONPRC18 Multimodal Macaque MRI atlas (anatomical MRI and diffusion MRI) - providing an anatomically comprehensive set of regions for the study of the Macaque brain [17]. Selected ROIs include the key regions affected in HD, such as regions in the basal ganglia circuit, cerebellum, prefrontal cortex, hippocampus, and amygdala (Table 1).

**Table 1.**
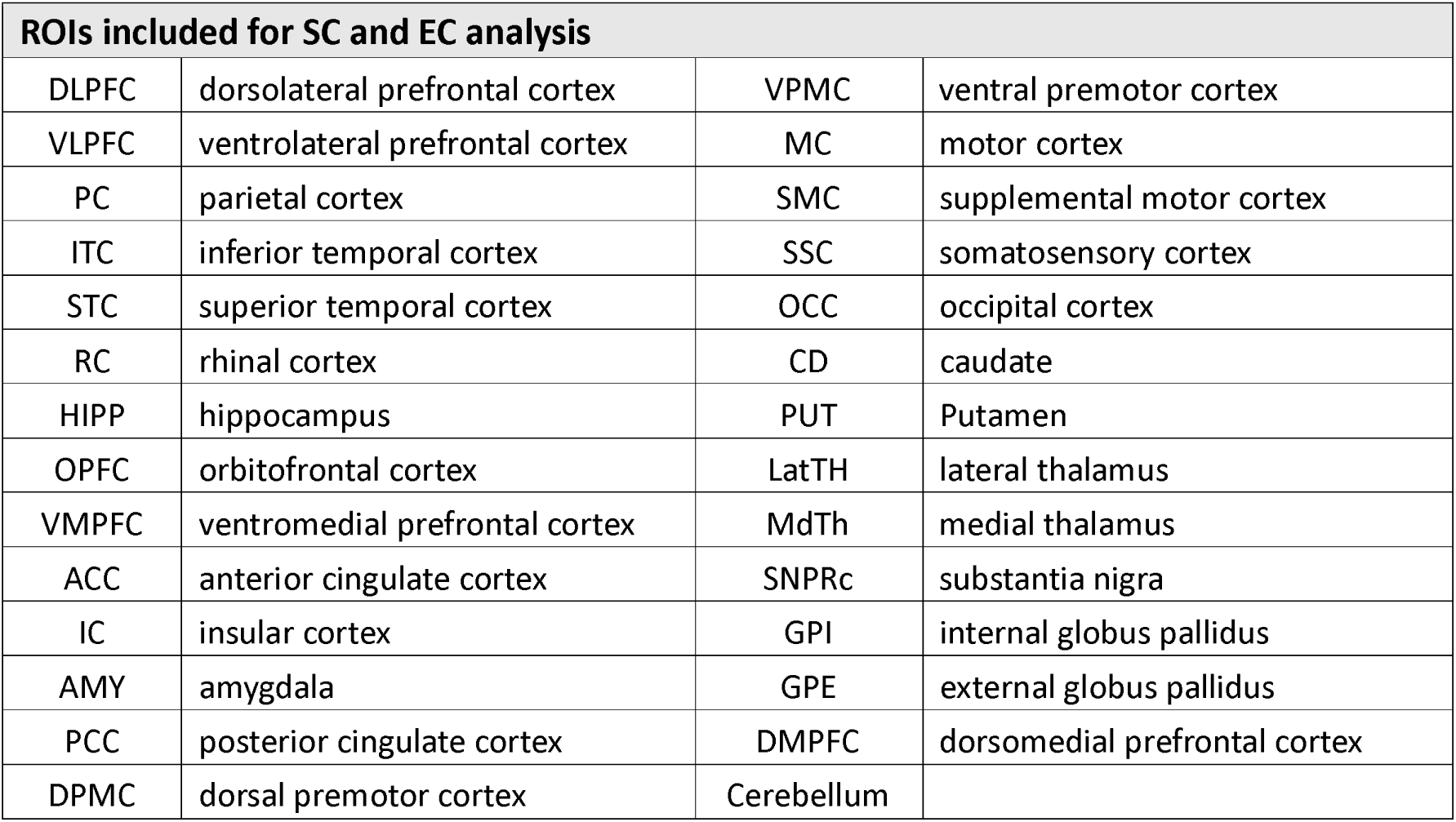
ROIs included for SC and EC analysis. The table lists the 28 ROIs used in this study to estimate SC and EC, along with their abbreviations. All ROIs, except for the cerebellum, are separated by hemisphere.

### 2.4 Structural Connectivity Estimation

Structural Connectivity estimation was carried out for each animal at each timepoint using MRtrix3.0 software [18] and in-house MATLAB code. For each subject and timepoint, the diffusion tensor was estimated from the diffusion-weighted data in subject space and the fractional anisotropy (FA) map was derived [18, 19]. A tensor-based probabilistic tractography algorithm (Tensor_Prob, MRTrix) was used to calculate 2000000 streamlines [20] – a streamline follows the orientation of the principal eigenvector of the tensor and incorporates the following parameters constraints: the maximum angle in between successive steps (45°), minimum (5mm) and maximum (1000mm) length of tracks, and minimum tensor FA cutoff value (FA > 0.15). Afterwards, tractography maps were labelled using the ONPRC18 atlas [17] with 55 ROIs via ANTs software [21, 22]. The FA map of each subject at each time point was co-registered to the FA map of the ONPRC18 atlas [17]. The estimated warp fields were used to back transform ROI labels from the ONPRC18 atlas to subject space. Finally, for each animal, a structural connectome was generated, weighted by the number of streamlines (NOS) that connect each pair of ROIs, as reconstructed from the tractogram [23] (Figure 1A).

**Figure 1.**
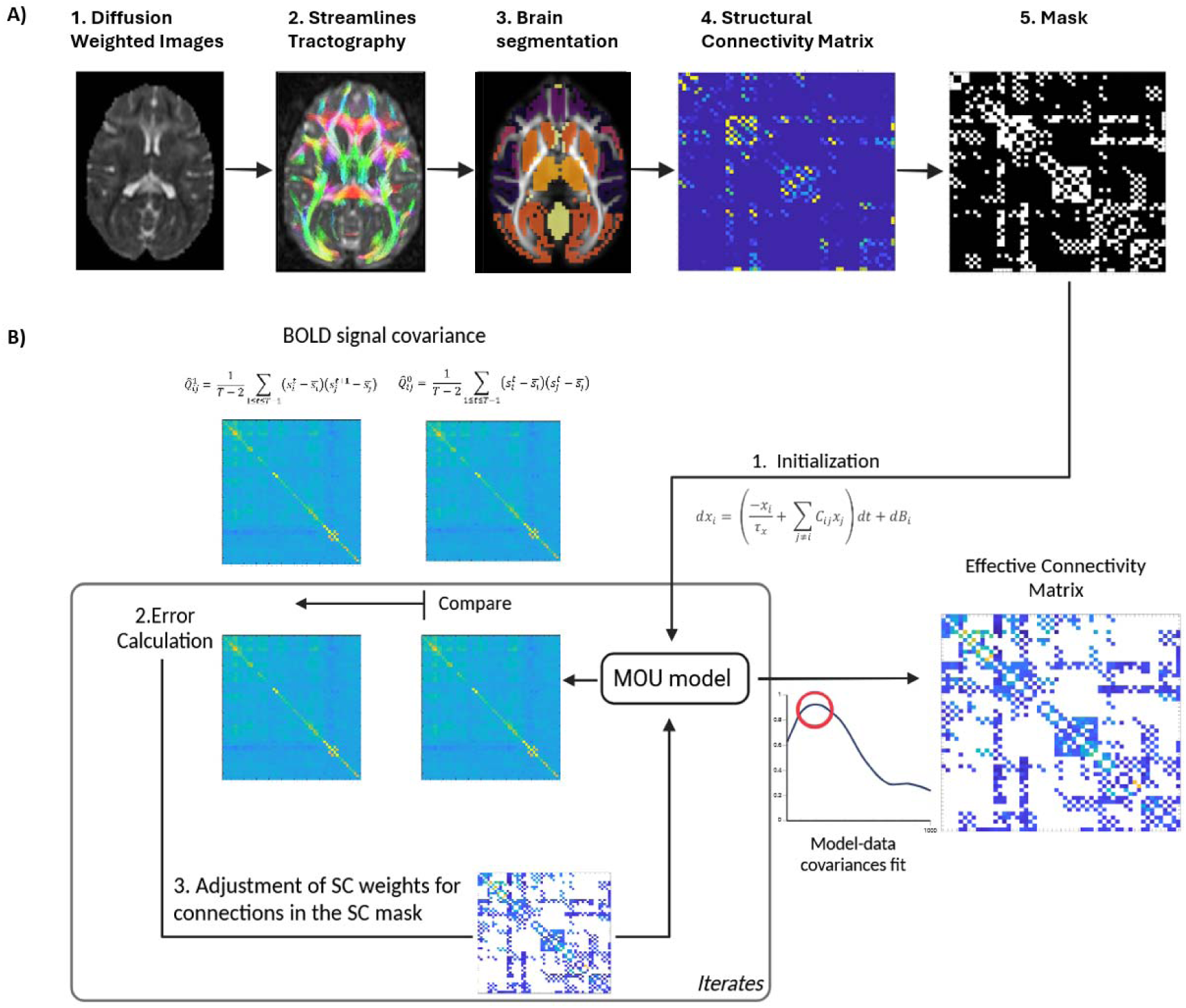
A graphical representation of the step-by-step process of the study. A) Structural Connectivity calculation: 1. The diffusion tensor was estimated from the diffusion-weighted data and the fractional anisotropy (FA) map was derived. 2. A tensor-based probabilistic tractography algorithm was used to calculate the streamlines. 3. Tractography maps were labelled. 4. A structural connectome was generated. 5. Using SC generated in 4, a binary mask is created for each subject consisting of top 25% strongest connections. B) Effective connectivity () estimation using a gradient descent algorithm: 1. Using the initial binary values the model equations are simulated and model covariances are calculated. 2. They are compared to empirical covariances and - zero and one-TR lagged covariances obtained from rs-fMRI BOLD signals and an error is calculated. 3. are adjusted based on the error and the process is repeated. The matrix of values that give the smallest error corresponding to the maximum Pearson correlation coefficient (marked with a red circle) between the model and empirical covariances for all connections (not only those included in the SC mask), is taken as effective connectivity.

### 2.5 Effective Connectivity Estimation

Effective Connectivity between the ROIs was estimated using a recently developed method which employs a multivariate Ornstein-Uhlenbeck (MOU) process model [13, 14] (Figure 1B).

First, from rs-fMRI data of each animal at each timepoint, zero-time lag and lagged (lag = 1TR) covariances between BOLD signals of ROIs were derived and used as empirical data in the model optimization step:

#### Estimation of lagged and zero lagged covariances

For each rs-fMRI scan comprising T time frames (index t), the BOLD time series are denoted by 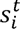 for every ROI (index i), where 1≤, *i* ≤ 55, and the mean ROI signal is denoted as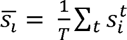. The For each rs-fMRI scan comprising T time frames (index t), the BOLD time series are denoted by t for empirical BOLD covariances without (1) and with (2) a time lag, typically 1 TR (∼ 2s, based on human studies), are subsequently expressed as follows:

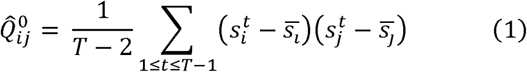

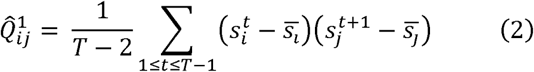

Then the structural mask, which serves as a prior for the strongest and biologically relevant connections, was generated for each subject at each time point. Towards this purpose, we used streamlines density, calculated by taking the number of streamlines between every pair of ROIs and dividing them by the sum of voxels comprising the two ROIs, as a measure of structural connectivity [18] and included 25% of strongest connections in the binary mask. The selection of a threshold of 25% was tailored to maintain a minimal connection density within the mask ensuring a robust fit between the model and empirical covariances for each individual animal while preventing overfitting [13, 16]. Though the connections in the basal ganglia circuit did not show streamline density above the threshold, these connections were still added because of their biological relevance in HD as well as in this model. Afterwards, MOU model optimization was performed to estimate EC for all connections included in the mask, that yielded an optimal fit between model and empirical covariances for all 55×55 connections for each subject at each time point.

#### MOU process to model whole-brain dynamics

The variable representing neuronal activity, denoted as *x_i_*, for region *i*, undergoes exponential decay characterized by a time constant, τ*_x_*, and evolves contingent upon the activities of other regions:

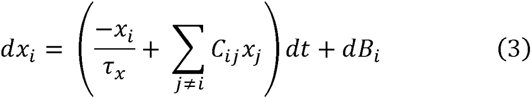

Here, τ*_x_* is associated with the exponential decay of the autocovariance averaged across all ROIs, the model, all variables, *dB_i_* are independent and correspond to a diagonal covariance matrix Σ. In the model, all variables, *x_i,_* have zero mean. *C_ij_* is connectivity, from region *j* to region *i*, which initially has a value of one or zero depending on whether the connection between these two regions is included in the mask. For each iteration connectivity is updated only for values which initially were set to one (included in the mask). The spatio-temporal zero lagged and lagged covariances are denoted by 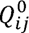 and 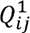 and can be calculated by solving the consistency equations below [13]:

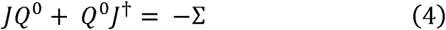

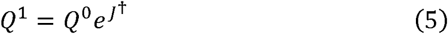

Here *J* is the Jacobian of the dynamical system; 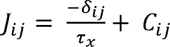, where Δ_*ij*_ is the Kronecker delta function and † denotes matrix transpose.

The final step is the optimization process, for which the objective is to adjust the parameters *C_ij_* of the model in such a manner that the covariance matrices *Q*^0^ and *Q*^1^ of the model, including all connections (not only those in the mask), replicate the corresponding empirical covariance matrices 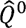 and 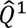. The connectivity update is:

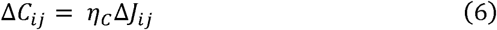

Here η*C* refers to the rate of change of connectivity which is set to 1E-4, in line with previous EC studies. The best fit corresponding to the minimum of error between the model and empirical covariances renders the optimal connectivity *C_ij_* that is then referred to as estimated EC.

### 2.6 Statistical Analysis

SC and EC values were statistically analysed using repeated measures ANOVA to assess the main effects of group (Buffer and 85Q), time (baseline, 3-, 6-, 9-, 14-, and 20-months), and group x time interaction. 30-month data was not included as data were missing for one animal.

The biodistribution of AAV2.retro vector differed in different regions of the brain [6, 24].Hence, only ROIs exhibiting viral vector propagation [6] were included in the statistical analysis for SC, resulting in the selection of 46 ROIs. For the EC statistical analysis, only ROIs with high viral vector propagation were considered, leading to the selection of 26 ROIs.

In case of statistically significant interaction effect (p < 0.05) was found after repeated measures ANOVA for a specific connection (either structural or effective), post-hoc group comparisons were performed using the least significant difference (LSD) test along with false discovery rate (FDR) for multiple post-hoc comparisons and all connections showing significant interaction effect. If no significant interaction was observed, only main effects were assessed followed by appropriate post-hoc comparisons and corrections for multiple comparisons.

We also carried out SC and EC statistical analysis of the three groups (Buffer, 10Q, and 85Q) in the same way as described above, for completeness and to relate to previous studies. The results are described in the supplementary materials (Supplementary Figures 6 and 7).

## 3. Results

### 3.1 Structural Connectivity Alterations

The design of the study by Weiss et al. [6] included three groups – the Buffer group receiving a buffered saline injection, a 10Q group overexpressing a 10 CAG repeats fragment of the human *HTT* gene and an 85Q group with a pathogenic 85 CAG repeats fragment of human *HTT* gene. In this work, we primarily focused on assessing the SC and EC changes in the 85Q group compared to the Buffer group for several considerations. First, even though we confirmed that the overexpression of the 10 CAG repeats fragment did not lead to the formation of aggregates in the 10Q group [8], the 10Q showed significant differences in working memory and motor deficits at multiple time points post-surgery not only with the 85Q group but also the Buffer group [6]. Additionally, we observed a diverging profile in brain metabolism towards a partly higher glucose uptake in the 10Q group which might be driven by imbalance in neuronal susceptibility due to the presence of excess 10Q protein [7]. Collectively, those findings might indicate a potential influence of the 10 CAG repeats overexpression on brain homeostasis. Therefore, we primarily focused on assessing the SC and EC changes in the 85Q group compared to the Buffer group.

We first derived mean SC for each group (Buffer, 85Q) at baseline, 3-, 6-, 9-, 14-, and 20-months post virus injection surgery and represented it as a circular graph depicting 46 ROIs from both hemispheres selected for statistical analysis. Figure 2 displays the SCs for the Buffer and 85Q group at the baseline (Figure 2A), and at timepoints when significant difference between the groups was found, namely, at 14 (Figure 2B), and 20 months (Figure 2C) post-surgery. The color-coded lines connecting two ROIs represent the connection strength using three different line weights, each representing a distinct category: thin lines indicate weak connections (from 1000 to 3000 streamlines), medium lines denote moderate connections (between 3000 and 7000 streamlines), and thick lines represent strong connections (over 7000 streamlines) between the two ROIs.

**Figure 2.**
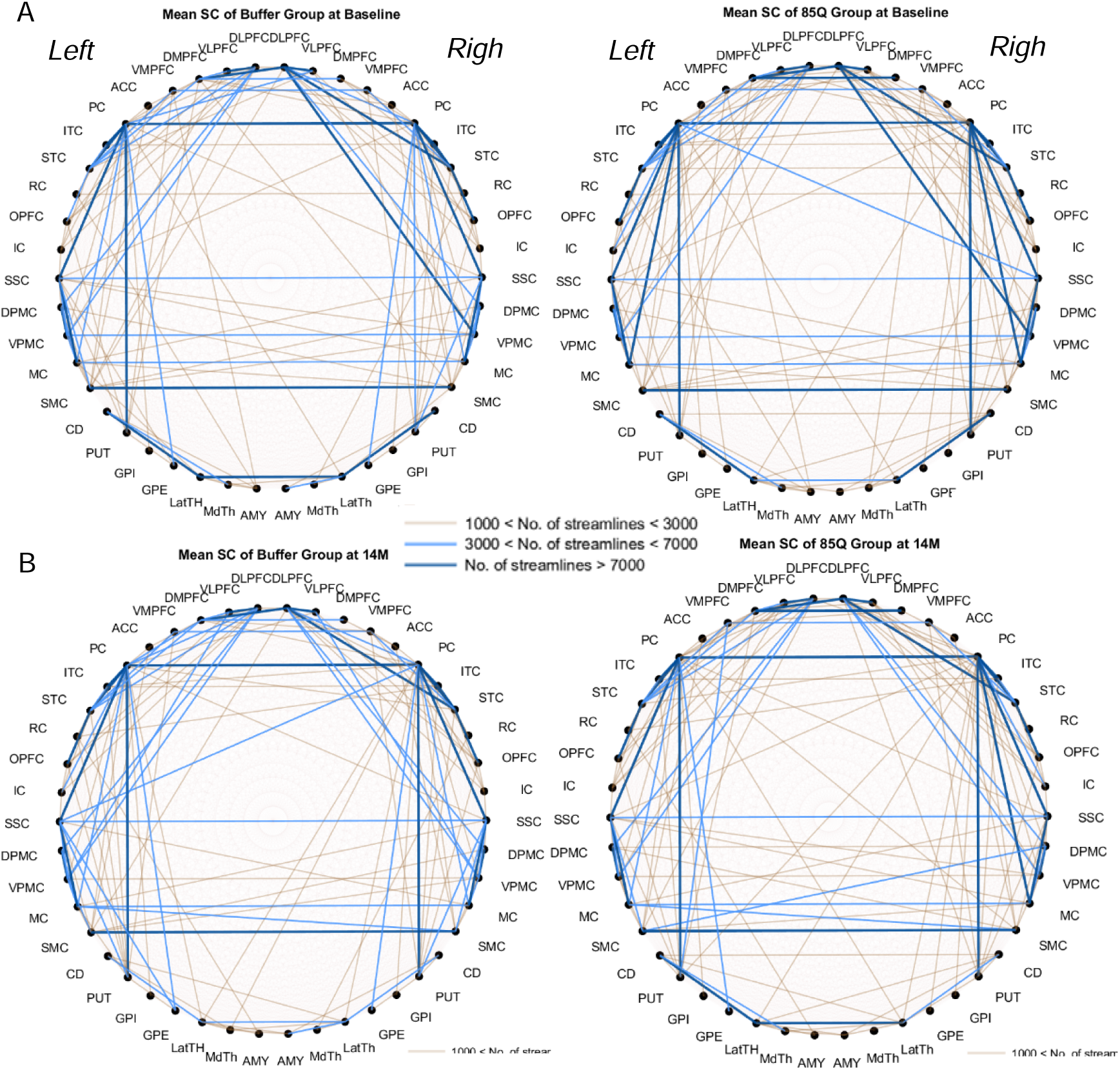
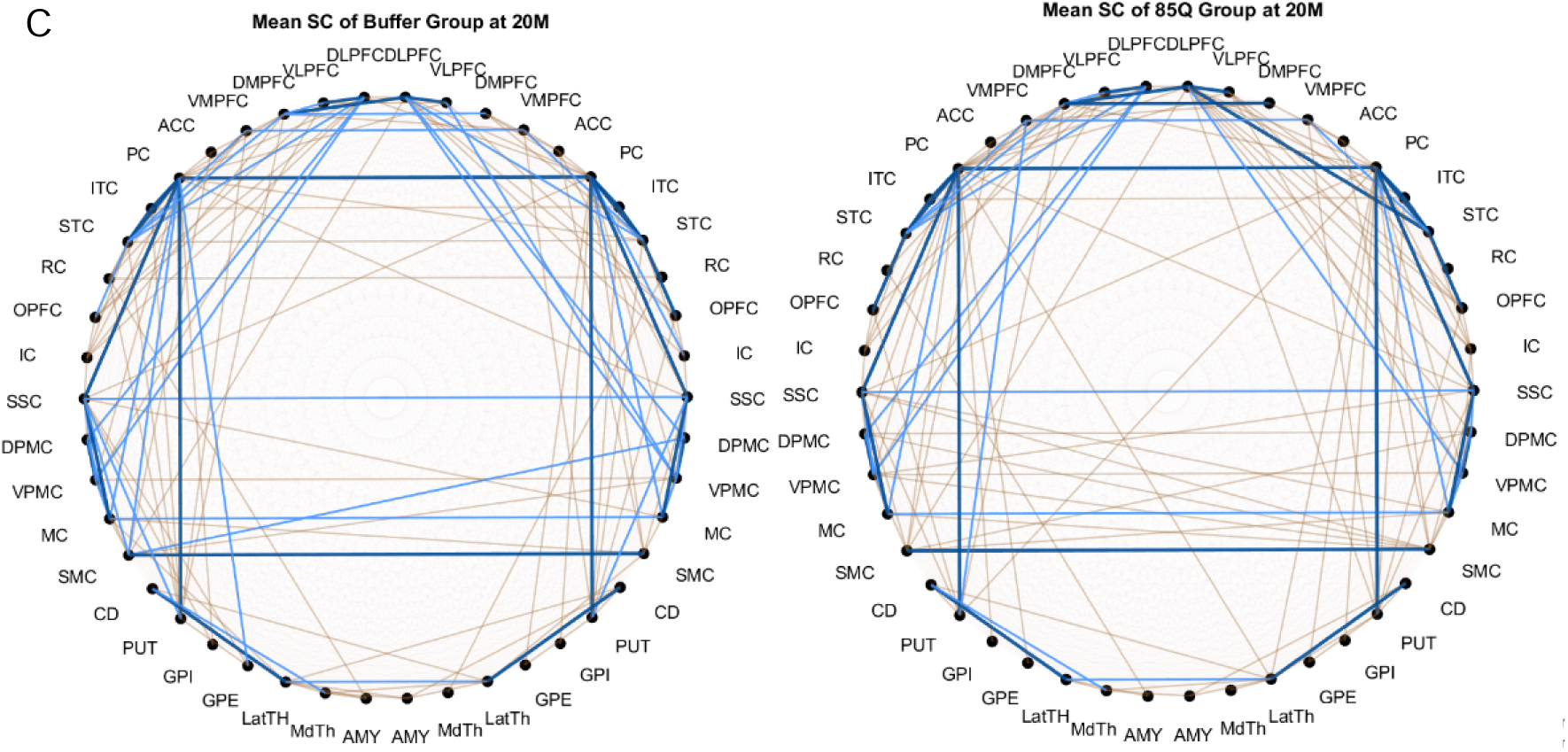
Mean SC in the Buffer and 85Q groups at baseline, 14 months, and 20 months. A circular diagram illustrates 46 ROIs divided into left and right hemispheres. Connection strength between ROIs is represented by line color: dark blue indicates more than 7000 streamlines, light blue indicates 3000–7000 streamlines, and yellow indicates fewer than 3000 streamlines.

Mean SCs of both groups at the baseline, and at 14 and 20-month timepoints showed that majority of the strong connections (> 7000 streamlines) were between the cortical regions. These include intrahemispheric connections such as those between the ventrolateral prefrontal cortex and dorsolateral prefrontal cortex, as well as between the parietal cortex and regions like inferior temporal cortex, superior temporal cortex, and somatosensory cortex. Additionally, we observed strong interhemispheric connectivity between the regions of the motor cortex (Figure 2 A-C). Supplementary Figure 1 shows the mean SC for the Buffer and 85Q groups at 3-, 6-, and 9-month time points.

We then assessed the group, time, and group x time interaction effect on SC between the 46 regions-of interest shown in Figure 2. Several connections between 42 of the 46 ROIs we considered showed a significant interaction effect (p < 0.05, repeated-measures ANOVA) as shown in either blue or red color in Figure 3A-B or Figure 4A-B. Interestingly, after post-hoc comparisons, none of these connections showed any significant inter-group differences at baseline, or at 3-, 6-, or 9-month timepoints. However, at the 14-, and 20-month timepoints we did observe significant difference between the Buffer and the 85Q group for some connections (Figure 3A, 3B, asterisks). Specifically, at the 14-month time point, we observed that the connection strength of global pallidus with putamen, somatosensory cortex, and parietal cortex, and of inferior temporal cortex with globus pallidus internal within the left hemisphere, as well as between inferior temporal cortex and superior temporal cortex in the right hemisphere was significantly lower in the 85Q group compared to the Buffer group (Figure 3A, C). At 20 months post-surgery, compared to the Buffer group, the 85Q group showed lower connection strength between rhinal cortex and putamen, somatosensory cortex and globus pallidus internal within the left hemisphere as well as between inferior temporal cortex and superior temporal cortex in the right hemisphere (Figure 3B-C).

**Figure 3.**
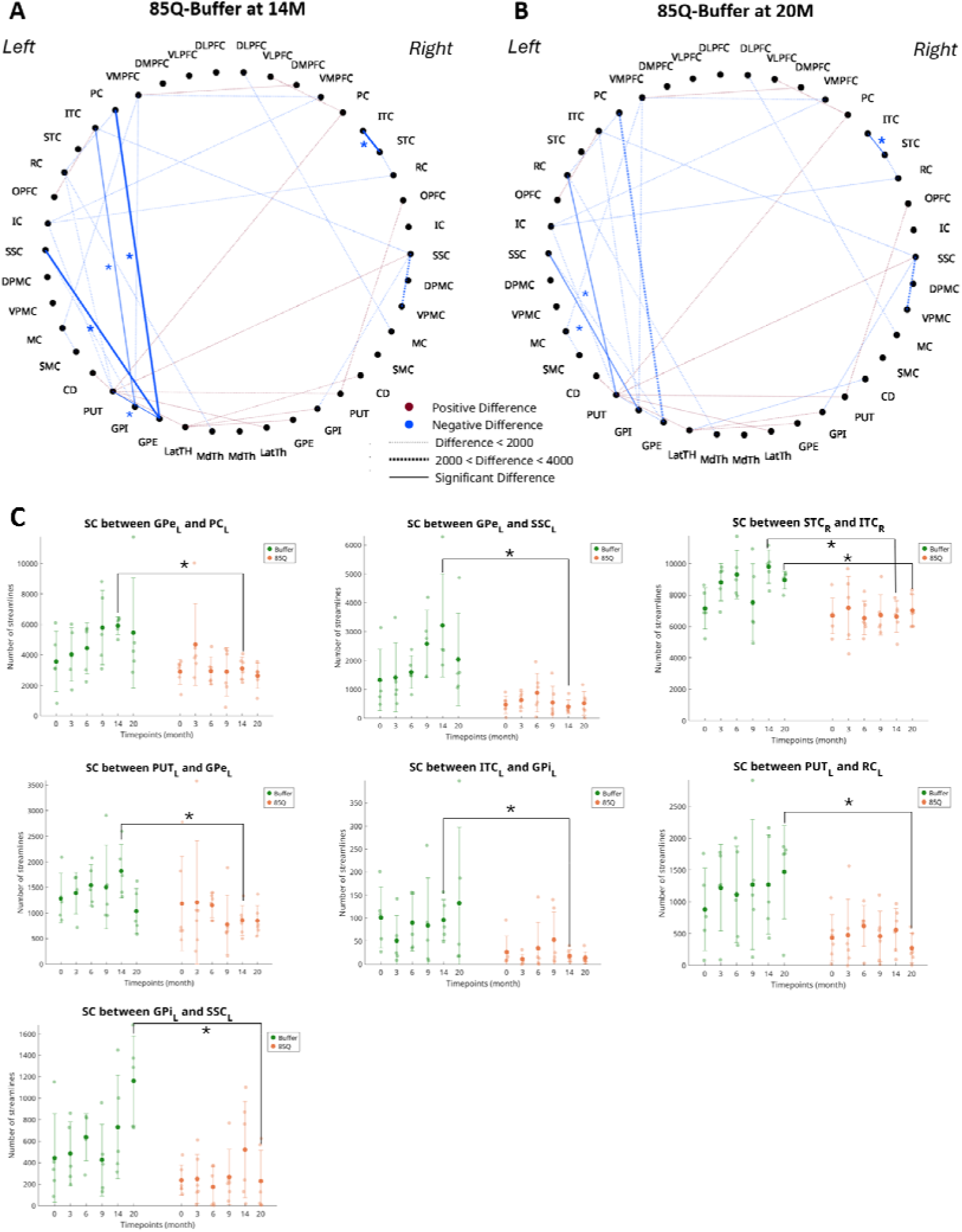
SC comparisons between 85Q and Buffer groups. A, B: A circular diagram illustrating connections (dashed lines) between 42 ROIs where a significant group x time interaction effect was found on SC and those (solid lines with asterisks) that showed significant difference on post-hoc comparisons between 85Q and Buffer groups at 14 months (A) and at 20 months (B). Blue/red colour indicates a lower/higher SC in 85Q as compared to Buffer. C) Mean +/-SD values of SC for each group at six timepoints between the ROIs where the post-hoc analysis showed significant inter-group difference either at 14 months or at 20 months after the post-hoc analysis.

**Figure 4.**
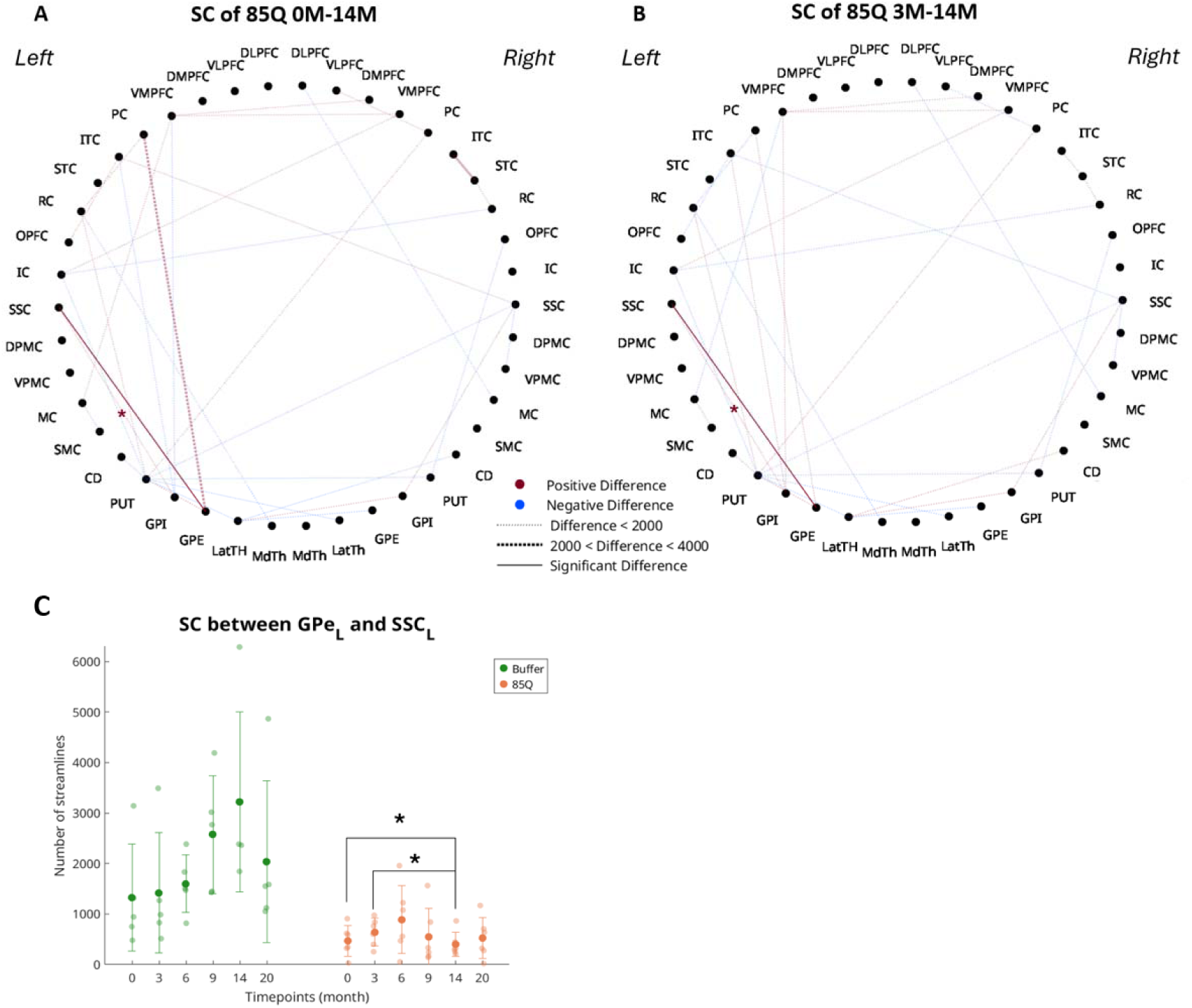
SC differences within 85Q group between different timepoints. A) A circular diagram illustrating all connections between 42 ROIs where a significant group x time interaction effect (p<0.05, repeated measures 2-way ANOVA) was found on SC between ROIs within 85Q group between 0 and 14 months B) A circular diagram illustrating 40 ROIs where a significant interaction effect was found in streamline count (SC) differences between ROIs within 85Q between 3 and 14 months. Dashed line indicates a significant ANOVA effect while solid line with an asterisk indicates a significant difference after post-hoc analysis. Blue/red color indicates a lower/higher SC in 85Q as compared to Buffer. C) Discrete scatter graph showing mean SC and standard deviation of both 85Q and control groups at six timepoints between the ROIs where the difference remained significant after the post-hoc analysis. A significant decrease in SC between globus pallidus external and somatosensory cortex in 85Q group at 14 months compared to 0 and 3 months is demonstrated.

Additionally, we found significant SC differences within the 85Q group between different timepoints. Connection strength between globus pallidus external and somatosensory cortex within the left hemisphere was significantly lower at 14 months compared to the baseline and at 3 months post-surgery (Figure 4).

Multiple significant group differences were observed between 85Q and Buffer groups (Figure 5), indicating that, irrespective of time, the two groups exhibited differences in interhemispheric SC between several regions: the anterior cingulate cortex and motor cortex, the parietal cortex and rhinal cortex, the amygdala and superior temporal cortex, and between left and right caudate nuclei. Additionally, a significant group difference was found in SC within the right hemisphere between the parietal cortex and rhinal cortex.

**Figure 5.**
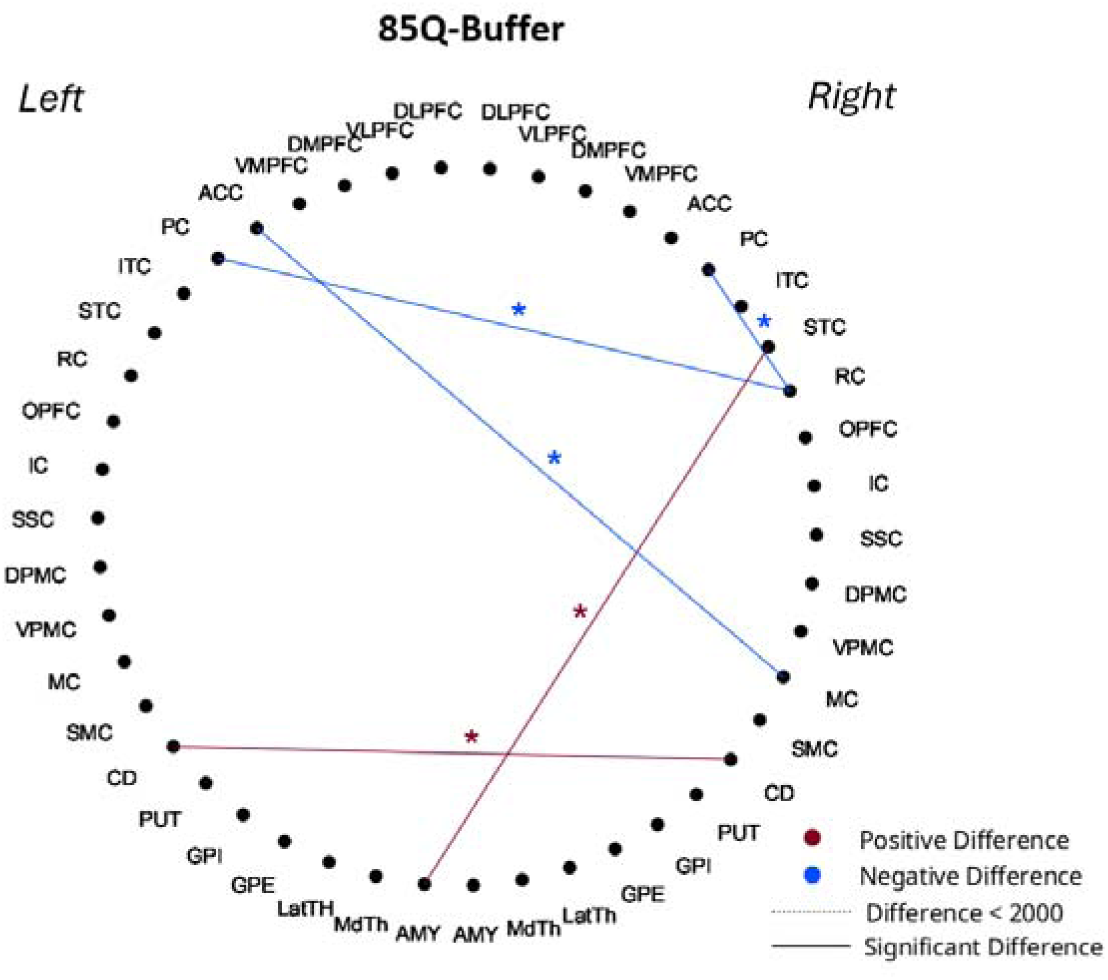
Group effect on SC values. A circular diagram illustrating connections between 46 ROIs that showed a significant main effect of group on SC (p < 0.05, repeated measures 2-way ANOVA). Blue/red color indicates a lower/higher SC in 85Q as compared to Buffer.

The ANOVA p-values, degrees of freedom and F-statistics for all structural connections that showed either a significant interaction or group effect and showed significant differences in post-hoc comparisons are described in Supplementary Table 1.

Supplementary Figure 2 shows the mean SC for the 10Q group at all six timepoints. When the 10Q group was included in statistical comparisons, we found that, in the 85Q animals, SC of GPE with the PC and SSC within the left hemisphere and the right hemispheric ITC-STC SC at 14-months post-surgery as well as left hemispheric SSC-GPI and RC-PUT SC at 20-months post-surgery remained significantly reduced when compared to the Buffer group (Supplementary Figure 6A-B). The 10Q group showed a significantly lower left hemispheric ITC-RC SC and right hemispheric ITC-STC SC compared to the 85Q group (Supplementary Figure 6C) and the Buffer group (Supplementary Figure 6D), respectively, at 14-months post-surgery. Additionally, at 20-months post-surgery, the 10Q group showed significantly reduced right hemispheric ITC-STC SC, left hemispheric RC-PUT and SSC-GPI SC, as well as an increased left hemispheric SC between ACC and SSC, compared to the Buffer group (Supplementary Figure 6E). The SC between SSC and DMPFC within the left hemisphere in the 10Q group was reduced at 20-months compared to 3-months post-surgery (Supplementary Figure 6F).

#### Effective Connectivity Alterations

Next, we estimated the individual subject’s EC for all connections included in the individual’s SC mask. The model optimization was assessed by calculating the maximum Pearson correlation coefficients between empirical covariances and MOU model generated covariances for each subject at each time point. These model fit values were 0.87 +/-0.06 SD for the Buffer group, 0.9 +/-0.07 SD for the 85Q group (Supplementary Figure 3). Supplementary Figures 4 and 5 show the group averaged EC matrices at baseline, 3-, 6-, 9-, 14-, and 20-months post-surgery for the Buffer and 85Q group respectively. The asymmetry of EC values reflects the directional information captured by EC.

We investigated the group, time, and group x time interaction effects on the EC values between 26 out of 55 ROIs selected based on their biological relevance in HD or the viral vector distribution in these animals. We found two connections – from caudate to supplementary motor cortex in the right hemisphere and from putamen to motor cortex in the left hemisphere - showed a significant interaction effect (p < 0.05, repeated measures 2-way ANOVA; blue arrows Figure 6A-B). Post-hoc comparisons revealed that the EC from caudate to supplementary motor cortex in the right hemisphere at 3-months (Figure 6A, 6C-left panel) while from putamen to motor cortex at 14-months post-surgery was significantly lower in the 85Q group compared to the Buffer group (Figure 6B, 6C-right panel).

**Figure 6.**
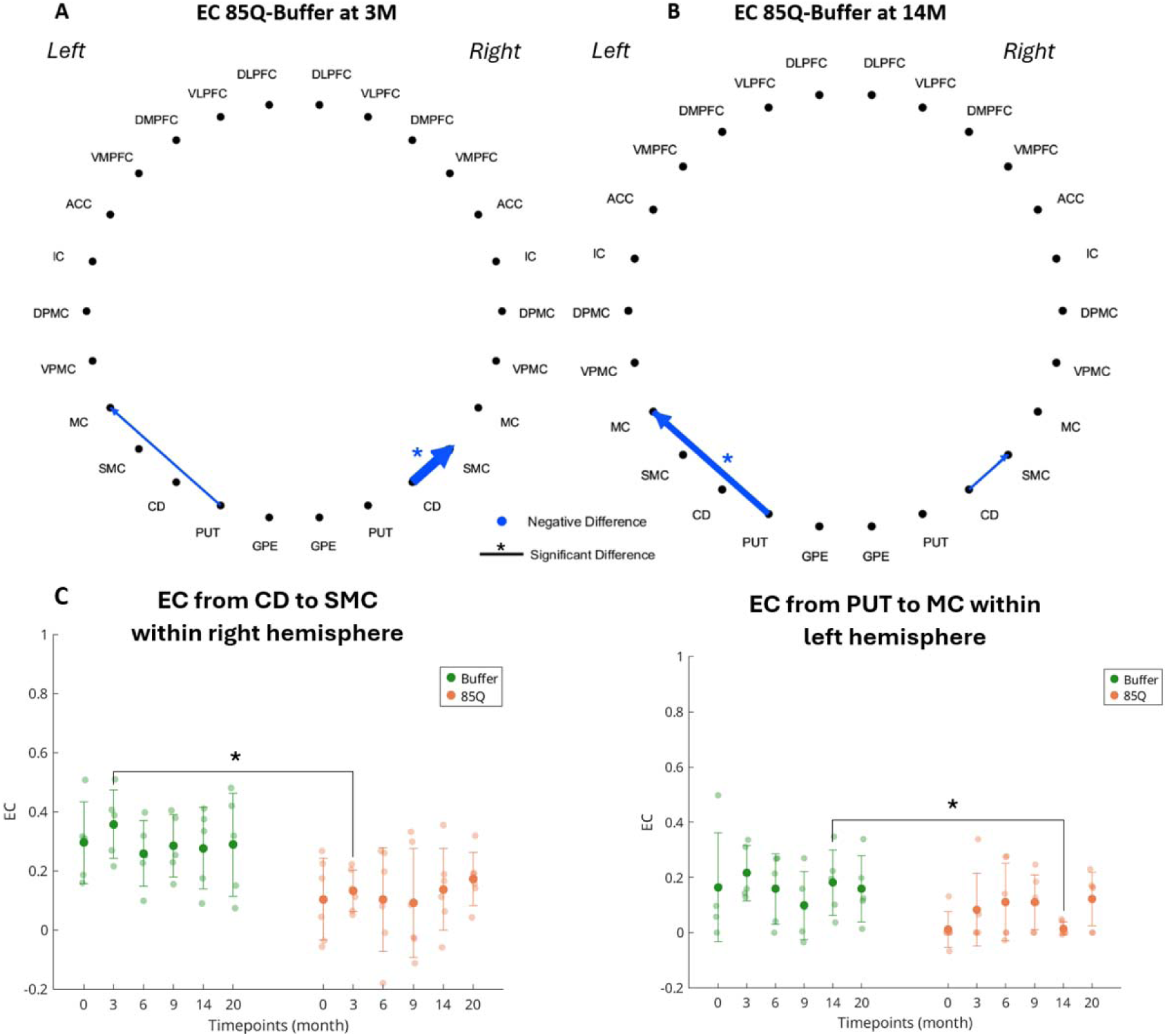
EC comparisons between 85Q and Buffer groups. *A, B:* A circular diagram illustrating connections (solid lines) between 26 ROIs where a significant group x time interaction effect was found on EC and those (solid lines with asterisks) that showed significant difference on post-hoc comparisons between 85Q and Buffer groups at 3 months (A) and at 14 months (B Blue/red color indicates a lower/higher EC in 85Q as compared to Buffer. C) Mean +/-SD values of EC for each group at six timepoints between the ROIs where the post-hoc analysis showed significant inter-group difference either at 3 months or at 14 months after the post-hoc analysis.

We also found a significant group effect for one connection; the EC from anterior cingulate cortex to ventral medial prefrontal cortex in the right hemisphere was significantly lower in the 85Q group compared to the Buffer group (Figure 7). The ANOVA p-values, degrees of freedom and F-statistics for all connections whose effective connectivity showed either a significant interaction or group effect and showed significant differences in post-hoc comparisons are described in Supplementary Table 2.

**Figure 7.**
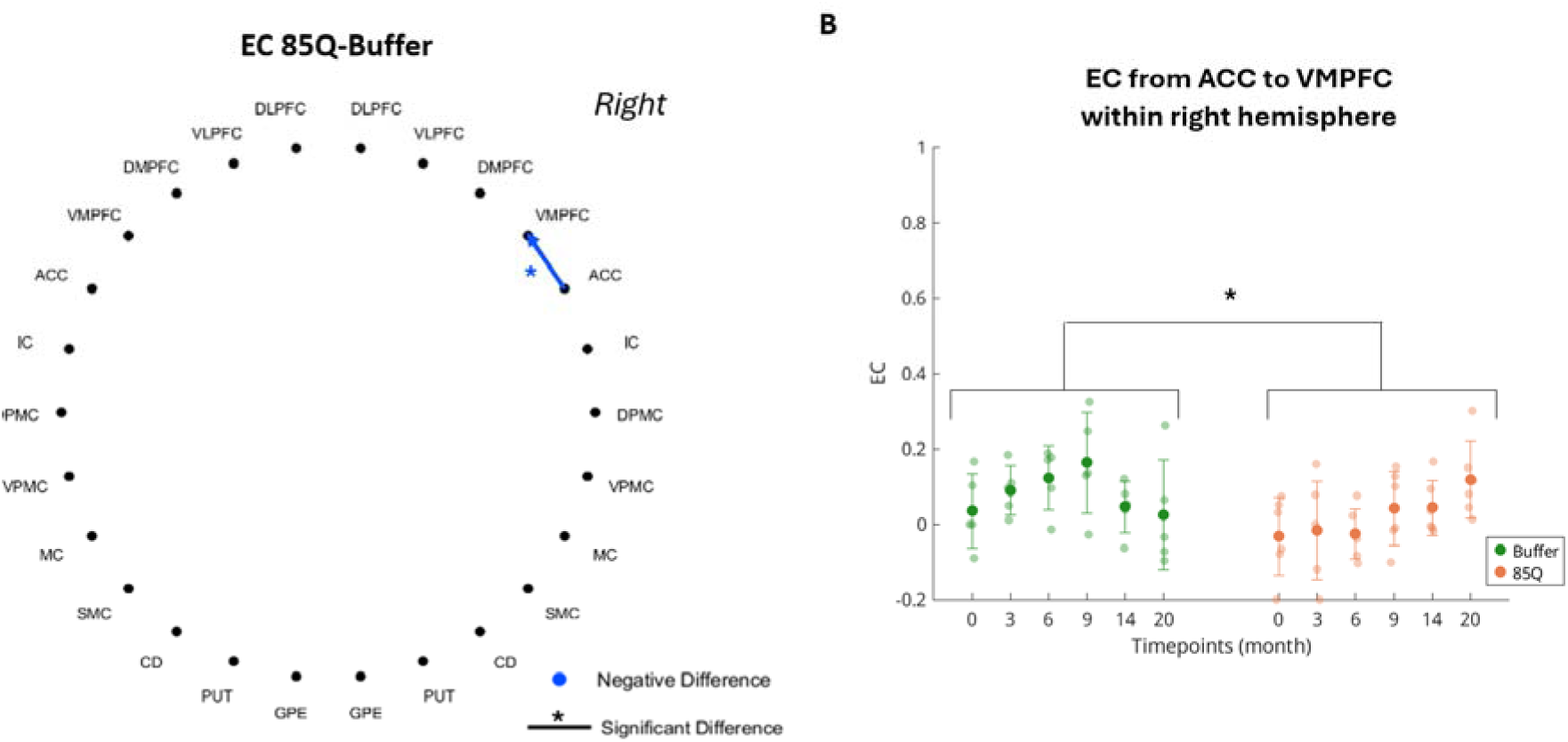
Group effect on EC values. A) A circular diagram illustrating difference in EC for connections between 36 ROIs that showed a significant main effect of group on EC (p < 0.05, repeated measures 2-way ANOVA). Blue line indicates a negative difference. B) Mean +/-SD values of EC for each group at six timepoints between the ROIs where the post-hoc analysis showed significant group difference after the post-hoc analysis.

When the 10Q group was included for statistical comparisons, the EC from the Caudate nucleus to Putamen in the right hemisphere was found to be higher in the 10Q group compared to the 85Q group at 3- and 9-months post-surgery (Supplementary Figure 7A-B). Further, the EC from dorsolateral prefrontal cortex to insular cortex in the right hemisphere was lower in the 10Q group compared to the 85Q group at 9-months post-surgery (Supplementary Figure 7B). Finally, the EC from the right anterior cingulate cortex to the left ventromedial prefrontal cortex showed a significant group effect and was found to be higher in the 10Q group compared to 85Q group irrespective of time post-surgery (Supplementary Figure 7C). No difference was found in the EC for any connection between the 10Q group and the Buffer.

## 4. Discussion

In this study, we aimed to assess longitudinal structural and effective connectivity changes in a recently developed NHP model of HD using AAV variants. In the 85Q model animals, the injection of the viral vector resulted in the expression of the vector leading to the production of the 85Q fragment in the caudate, putamen, and various other cortical and subcortical regions sending afferent projections to them [1,13]. Our previous PET studies detected mHTT aggregate formation in several brain structures and altered dopaminergic D2/3 receptor density and glucose metabolism in 85Q monkeys [7, 8]. In comparison with the Buffer group, 85Q animals also exhibited HD-typical behavioral readouts, mild tissue atrophy and microstructural alterations in striatal and cortical regions, and reduced rs-FC between sensory-motor areas and a rs network comprising of the striatum, ventral sensory motor cortex, and medial prefrontal cortex [6]. In our study, we found significant reductions in SC in connections involving multiple HD-associated regions in 85Q animals vis-à-vis the Buffer group at predominantly the latter timepoints post the virus injection surgery. On the other hand, a reduction in the EC from the caudate nucleus to the supplementary motor cortex in the 85Q group was detected at just 3-months post-surgery indicating early changes in causal, functional interaction in HD-relevant regions.

### 4.1 Structural Connectivity Changes

After generating a diffusion-tensor based tractography from the diffusion-weighted MRI, SC between pairs of grey matter regions was quantified as the number of streamlines within the tractogram, building a structural connectome. No significant differences in SC were found prior to the virus injection surgery between the groups, despite variations in age and sex between the animals.

The pathogenesis of HD originates in the striatum (comprising the caudate and putamen), where mHTT first impacts neurons bearing D2 receptors, leading to neurodegeneration, which, in turn, affects the afferent projections extending to GPE, subsequently, disrupting the entire indirect pathway in basal ganglia. In the 85Q animals, [F]Fallypride PET imaging showed reduced D2/D3 receptor density in the caudate, putamen and globus pallidus external compared to both the Buffer and 10Q groups [7]. In later stages of the disease, neurons of the direct D1-pathway are also affected. Additionally, as the disease progresses pathology extends to cortical regions, particularly the motor cortex [1]. Consequently, at the early stages of the disease, MRI techniques reveal volume loss and altered diffusion metrics within basal ganglia regions in PwHD [25]. In the 85Q NHP model of HD considered in this study, DTI showed reduced FA in caudate nucleus and motor and other cortices while increased FA in the putamen and globus pallidus from the early timepoint of 3-months post-surgery [9]. Additionally, studies in PwHD reported increased diffusivity and reduced FA in symptomatic HD in prefrontal WM tracts, corpus callosum, and cortico-striatal tract [26], along with altered connectivity between the putamen and prefrontal/motor cortices, later extending to caudate-parietal cortex connections [27].

Given the disease mechanism and previous volumetric and structural changes in this NHP model, we expected that SC changes would initially appear within the basal ganglia, then extend to the motor cortex, and subsequently involve frontal and prefrontal regions in the 85Q. Instead, we found significantly reduced SC between the basal ganglia regions (such as the putamen, globus pallidus internal, and globus pallidus external) and the cortex, as well as between a few cortical areas at 14- and 20-months post-surgery. These findings suggest that the 85Q animal model partially recapitulates key features of the HD phenotype. The explanation for such results can be that this NHP model of HD is developed by surgically injecting a viral vector, hence mHTT is expressed in specific regions, rather than throughout the whole brain. In a prior study on a different cohort, the expression of mHTT following intra-caudate and intra-putamen injection of AAV2.retro-HTT85Q was assessed in multiple regions. Brain regions were qualitatively ranked relative to one another, considering cell number and density, and placed into categories of high, medium, low, and minimal/none [6, 24]. Most of the regions known to be affected in the early stages of HD such as GPI, SNPRC, MdTh, LaTh showed low expression levels of mHTT [6, 24]. Thus, it is possible that the concentration of mHTT within these regions was insufficient to induce any detectable changes in SC within basal ganglia. Despite this, we found reduced SC between left GPI and left ITC (low mHTT expression) at the 14-month timepoint and between left GPI and left SSC (medium mHTT expression) at the 20-month timepoint. On the other hand, the putamen (PUT), which served as an injection site for the viral vector, and globus pallidus external (GPE), showed high and medium levels of mHTT expressions respectively. Consequently, multiple structural connections involving these regions (such as PUT-RC, GPE-PC, GPE-PUT, GPE-SSC) were reduced in the 85Q group compared to Buffer.

Additionally, the [^11^C]CHDI-180R PET imaging study showed that in the 85Q NHP model of HD utilized in this study, the expression of mHTT is significantly higher in regions such as DMPFC, ACC, SMC, and PUT compared to the Buffer and 10Q groups [13]. Some of these regions showed corresponding reductions in structural connectivity, such as (PUT-GPE, PUT-RC). However, we also observed SC reductions in connections such as the GPE-PC, GPI-SSC, and ITC-STC, where no significant mHTT expression was detected by PET. These findings suggest a more complex relationship between mHTT expression and structural connectivity changes in HD, implying that biological mechanisms other than mHTT accumulation may also contribute to the observed SC changes.

Another observation was that significant differences in SC were observed unilaterally, mostly in the left hemisphere. In PwHD unilateral atrophy or unilateral changes in SC have not been reported. In comparing our findings with those of Weiss et al. [6], it is important to highlight the different methodological approaches used in each study. Weiss et al. [6] observed minimal hemispheric asymmetry in their data and, consequently, conducted a voxel-wise analysis of FA across the entire brain without distinguishing between left and right hemispheres. In contrast, since our study utilized tractography to estimate number of streamlines between regions, we used regions in both hemispheres separately. This approach allowed us to detect significant reductions in SC specifically in the left hemisphere. However, this finding could have been influenced by the low statistical power of the study, which could limit the ability to detect significant differences consistently across both hemispheres.

For completeness in relation to previous studies in this model, we carried out statistical analysis comparing all three groups (Buffer, 10Q, and 85Q). However, the 10Q group displayed behavioural changes and motor symptoms similar to that of 85Q group, including significant differences in memory deficits, fine motor skill performance, and motor phenotypes at multiple timepoints [6]. In line with those previous observations, in this study we detected significant differences in SC when comparing the 1OQ group to Buffer as well as 85Q groups (Supplementary Figure 6). Majority of the significant SC differences we observed occurred when comparing Buffer and 10Q group at different timepoints (Supplementary Figure 6B-E) as well as within 10Q group between different timepoints (Supplementary Figure 6F). Collectively, this suggested that 10Q group resulted in partial deficits, although to a lesser extent than 85Q group, and hence, was not an optimal control group due to those intermediate effects.

### 4.2 Effective Connectivity Changes

Whole-brain structural connectivity has been shown to be related to functional connectivity especially during the resting-state [11, 12]. Resting-state functional connectivity alterations have been investigated in both PwHD [28] and in rodent models of HD [29, 30]. In PwHD, reduced FC has been found at manifest stages, but the findings are less consistent at early, pre-manifest stages. Rodent models of HD such as the zQ175 DN have shown a reduced cortico-striatal FC but only at 6 and 10 months of age [29]. Measures of temporal fluctuations in FC such as co-activation patterns (CAPs) and quasi-periodic patterns (QPPs) have been useful in identifying early changes [31, 32]. Specifically, we have shown hyperactivations in the default mode-like network regions in specific CAPs in 3-month-old zQ175DN mice before the manifestation of motor deficits observed at 6 months. In the 85Q NHP model of HD considered in our study, Weiss et al. found altered functional connectivity of several cortical and subcortical voxels with an independent component comprising of striatal and sensory-motor areas over the course of 30 months post-surgery [6]. However, all these studies investigate only correlative measures of statistical association between the neuronal activities of different brain regions indirectly measured by their BOLD signals. No prior study has assessed alterations in directed/causal functional influence from one region on to another in the context of HD.

Here, we investigated such directed functional interactions in the form of effective connectivity using a recently developed method in the 85Q NHP model of HD. MOU-EC method uses a binary, sparse mask of strong structural connections and finds optimal scaling weights associated with them so that simulated zero-lagged and lagged covariances between all regions match those observed empirically in the resting-state scan of individuals [13, 16]. These optimal weights, referred to as EC, give a measure of directed influence between regions. As we infer the EC by simulating a whole-brain network model, the explanation for observed values of lagged and non-lagged functional connectivity between any pair of regions optimally incorporates the importance of both direct and indirect structural pathways, via other regions, between them [33]. Also, as they are inferred from functional MRI data, the EC values reflect not just the density of axonal fibers measured by structural connectivity but also concentrations of receptors and neurotransmitters as well as local regional excitability [33].

EC has been shown to be more sensitive than standard, static FC as a distinguishing fingerprint of an individual based on their rsfMRI scan or in distinguishing healthy participants from stroke patients [16, 34]. In this study, we found a significantly reduced EC from the caudate to supplementary motor cortex within the right hemisphere in the 85Q group at three months post-surgery compared to the Buffer group. This aligns with the pathology of HD as the initial changes in PwHD primarily occur in the striatum, which in turn affects cortical areas through the indirect pathway [1]. We also observed a significant decrease in EC from putamen to motor cortex at 14 months within the left hemisphere in the 85Q group compared to the Buffer group – other key regions known to dysfunction in HD. No other EC differences within basal ganglia circuit were detected, but an overall significant reduction in EC from anterior cingulate cortex to ventromedial prefrontal cortex was observed in the 85Q group compared to the Buffer group irrespective of the time past surgery. All these regions showed a high propagation of viral vector and high expression of mHTT. In particular, the caudate, putamen, supplementary motor cortex and the anterior cingulate all exhibited significantly higher levels of binding of [^11^C]CHDI-180R tracer, implying significantly higher mHTT aggregates in these regions in the 85Q group compared with the Buffer group. Therefore, the reductions in EC in connections involving these regions may be related to the mHTT aggregation found within. Moreover, structural connectivity changes were not observed in these connections at any of these timepoints. This indicates that the changes in the EC of these connections could be related to the impact of mHTT aggregates on synaptic transmission rather than the axonal degeneration/microstructural changes in these fibers. This interpretation is supported by the findings of Weiss et al. [7], who reported significantly reduced D₂/D₃ receptor densities in the caudate and putamen – regions that, in the 85Q group, showed reduced EC to the motor cortex areas. Such reductions in receptor availability point to impaired dopaminergic neurotransmission, which could contribute to disrupted circuit dynamics. Furthermore, alterations in synaptic transmission have been reported in HD, supporting the characterization of the disease as a synaptopathy [35]. Consistent with this, previous studies have shown that disrupted synaptic function can lead to changes in FC [36]. Finally, the fact that reduced EC between the caudate and supplementary motor cortex was observed as early as 3-months post-surgery underscores the sensitivity of EC in detecting early changes.

### 4.3 Limitations of the study

One limitation of this NHP model of HD is that, unlike mouse models that are obtained using genetic mutations, here a viral vector carrying the fragment with 85 CAG repeats is injected in the caudate and putamen which then limits the expression and spread of mHTT to specific regions instead of systemically in the whole brain. Our analyses in this study, especially the estimation of EC is done at the whole-brain level to incorporate the influence of both direct and indirect pathways in the network. Therefore, the effect of the spread of mHTT expression only to specific regions may have contributed to diminished longitudinal alterations in EC which were indeed fewer than those in SC. Secondly, while the sample size is good for NHP studies, it may not have been sufficient for the analyses considered in this study especially at the level ROI-pairs and the number of repeated measures considered. As a result, the post-hoc analyses with multiple comparisons corrections may have been unable to consistently detect significant inter-group differences. Additionally, the age of subjects varied from 6 to 14 years old (prior the surgery), and groups were not sex matched. Weiss et al. observed that the capacity limitations of the viral vector packaging prevent the expression of full-length HTT gene, leading to an expression of only a fragment, which differs from wild type HTT and hence, can have an unexpected effect on the organism. Finally, the animal models of HD, differently to humans, retain their natural HTT genes, which may modulate the effects of the introduced mutant gene. This might explain the changes in behavior and the significant SC differences between 10Q, and Buffer group found in our study (Supplementary Figure 6).

In summary, our longitudinal investigation of structural and effective connectivity in the 85Q NHP model of HD revealed alterations in cortico-basal ganglia circuitry that partially recapitulate the phenotype of human HD. Structural connectivity analysis demonstrated significant reductions in connections involving the putamen and external globus pallidus primarily, consistent with localized mHTT expression. This pattern may reflect inherent susceptibility in these regions, potentially amplified by higher levels of vector expression. Interestingly, some SC changes occurred in regions with low or undetectable mHTT levels, indicating that secondary or network-level mechanisms may also contribute to connectivity disruption. Hierarchy in the topographic organization, especially of the cortical regions, that situates trans-modal regions like those from the default mode network on one end of the spectrum while unimodal ones like primary sensory and motor regions on the other has been reported in both humans and macaques [37]. In humans, graph theoretical analyses on structural connectivity obtained using DTI data have identified densely interconnected rich club regions that includes the putamen along with other default mode network regions [38]. Therefore, it is possible that transmodal and rich-club regions such as the anterior cingulate cortex and putamen, respectively, which show high levels of mHTT expression in this NHP model [8] could disrupt downstream structural pathways to other regions less impacted by the spread of mHTT alone. Future studies could use graph theoretical tools to study network effects in SC in this model as well as explore changes in cortical topographical hierarchy. Network effects in functional data were partly captured by our effective connectivity analysis that uncovered early differences in key striatal-cortical pathways and in regions that showed high mHTT aggregates. Together, these findings support the translational relevance of the 85Q macaque model and underscore the potential of multimodal connectivity metrics to capture the evolving pathophysiology of HD.

## Acknowledgments

This work was partly supported by National Institutes of Health grants NS099136, AG078407, OD011092 awarded to AW. The computational resources and services used in this work were provided by the HPC core facility CalcUA of the University of Antwerp, the VSC (Flemish Supercomputer Center), funded by the Hercules Foundation, and the Flemish Government department EWI.

## Data Availability statement

The datasets used and/or analyzed during the current study can be obtained by submitting a reasonable request to the corresponding author.

## 6. Supplementary figures

**Supplementary Figure 1.**
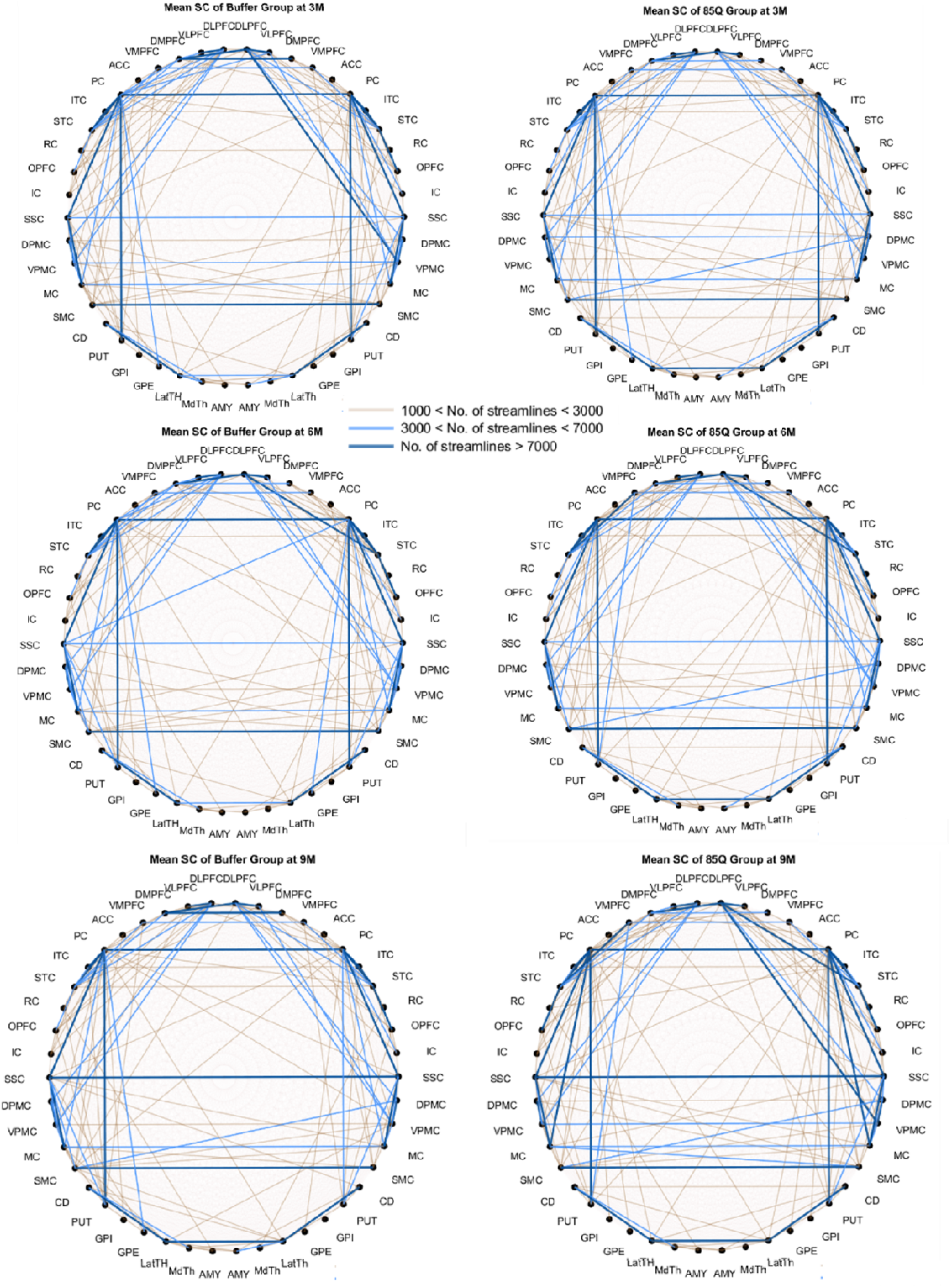
Mean SC in 85Q and Buffer groups at 3 months, 6 months, and 9 months. A circular diagram illustrates 46 ROIs divided into left and right hemispheres. Connection strength between ROIs is represented by line color: dark blue indicates more than 7000 streamlines, light blue indicates 3000–7000 streamlines, and yellow indicates fewer than 3000 streamlines.

**Supplementary Figure 2.**
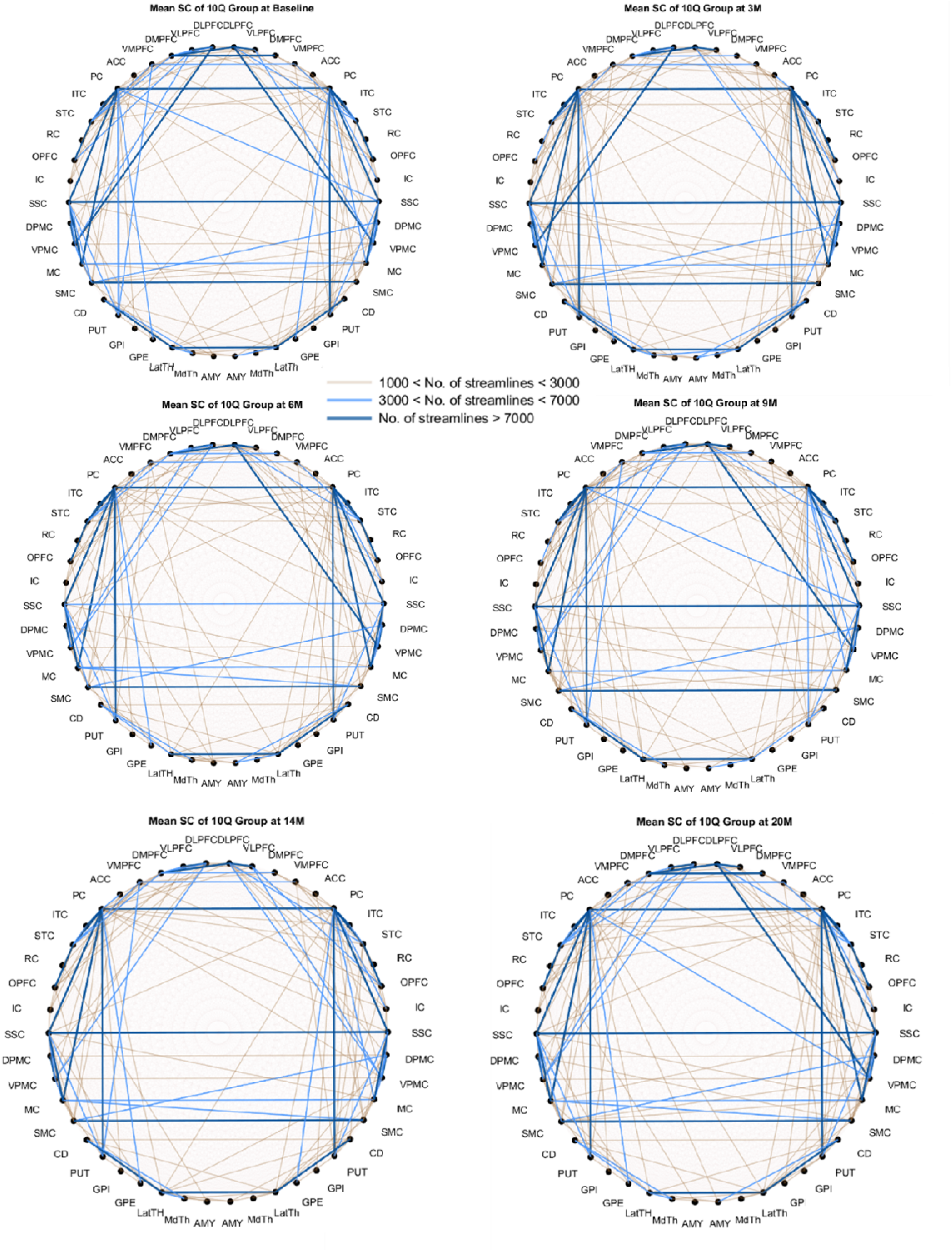
Mean SC in 10Q group at baseline, 3-, 6-, 9-, 14-, and 20-months A circular diagram illustrates 46 ROIs divided into left and right hemispheres. Connection strength between ROIs is represented by line color: dark blue indicates more than 7000 streamlines, light blue indicates 3000–7000 streamlines, and yellow indicates fewer than 3000 streamlines.

**Supplementary Figure 3.**
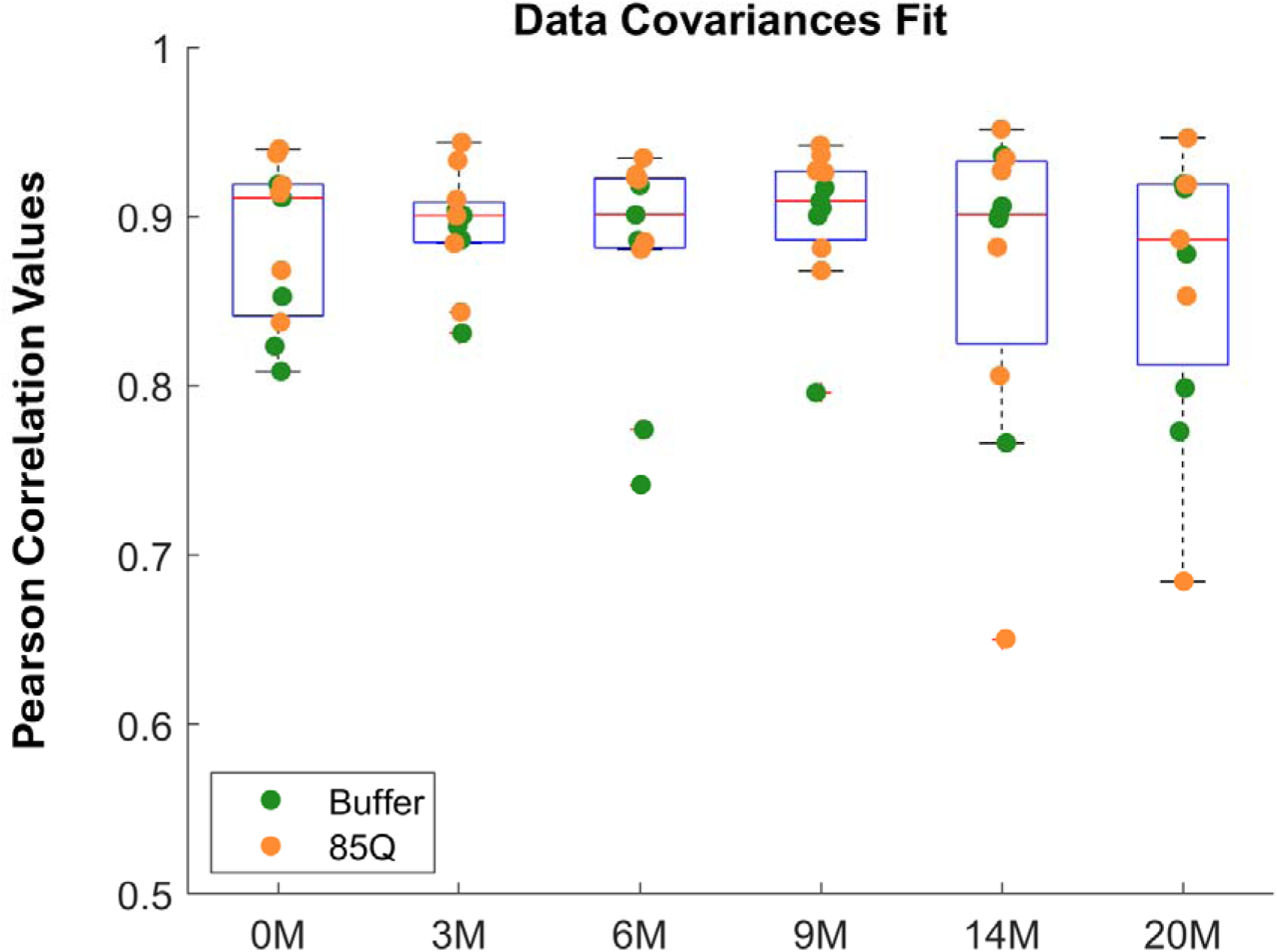
Max Pearson correlation values between empirical and MOU generated covariances for each subject at each time point.

**Supplementary Figure 4.**
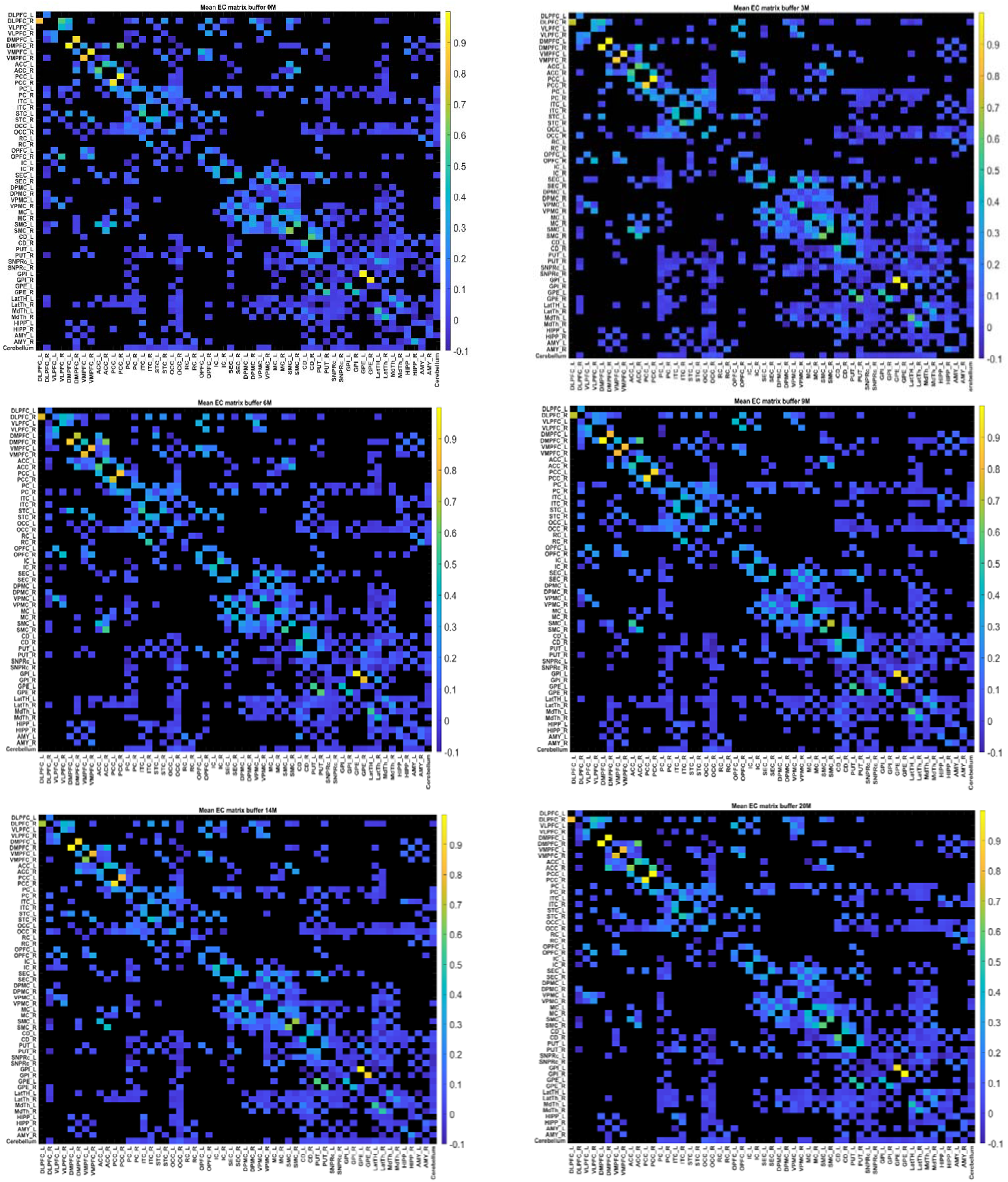
Mean effective connectivity in Buffer group at baseline, 3-, 6-, 9-, 14-, and 20-months. X and Y axes are ROIs.

**Supplementary Figure 5.**
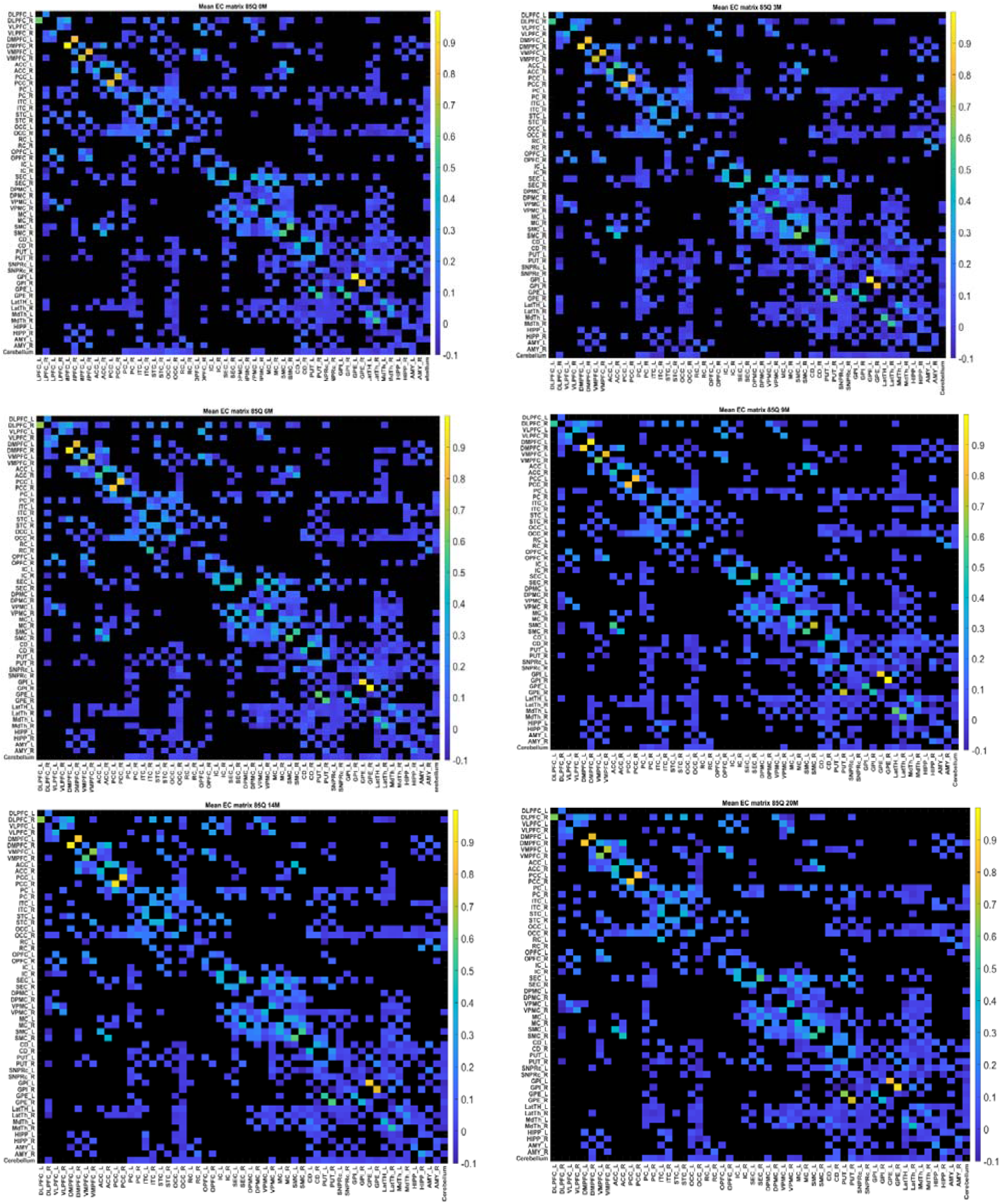
Mean effective connectivity in 85Q group at baseline, 3-, 6-, 9-, 14-, and 20-months. X and Y axes represent ROIs.

**Supplementary Table 1.** A table containing F-statistic, degrees of freedom and p-values for structural connections that showed a significant interaction or group effect followed by a significant difference after post-hoc comparisons.

| Connection Name | Effect Type | Degrees of Freedom | F statistic | P value |
| --- | --- | --- | --- | --- |
| GPe(L) – PC(L) | Interaction (Group x Time) | 1 | 4,56 | 0.0022 |
| GPe(L) – SSC(L) | Interaction (Group x Time) | 1 | 5.84 | 0.0259 |
| STC(R) – ITC(R) | Interaction (Group x Time) | 1 | 8.92 | 0.0112 |
| GPe(L) – PUT(L) | Interaction (Group x Time) | 1 | 4.68 | 0.0259 |
| ITC(L) – GPi(L) | Interaction (Group x Time) | 1 | 3.1 | 0.0259 |
| PUT(L) – RC(L) | Interaction (Group x Time) | 1 | 5.05 | 0.0474 |
| GPi(L) – SSC(L) | Interaction (Group x Time) | 1 | 9.36 | 0.0474 |
| STC(R)– ITC(R) | Interaction (Group x Time) | 1 | 8.915 | 0.0474 |
| Gpe(L)-SSC(L) | Interaction (Time x Group) | 1 | 5.84 | 0.0184 |
| Gpe(L)-SSC(L) | Interaction (Time x Group) | 1 | 5.84 | 0.0063 |
| RC(R)-PC(L) | Group | 1 | 3.39 | 0.0472 |
| RC(R)-PC(R) | Group | 1 | 4.18 | 0.0472 |
| STC(R)-AMY(L) | Group | 1 | 3.59 | 0.0472 |
| MC(R)-ACC(L) | Group | 1 | 4.2 | 0.0472 |
| CD(L)-CD(R) | Group | 1 | 3.14 | 0.0472 |

**Supplementary Figure 6.**
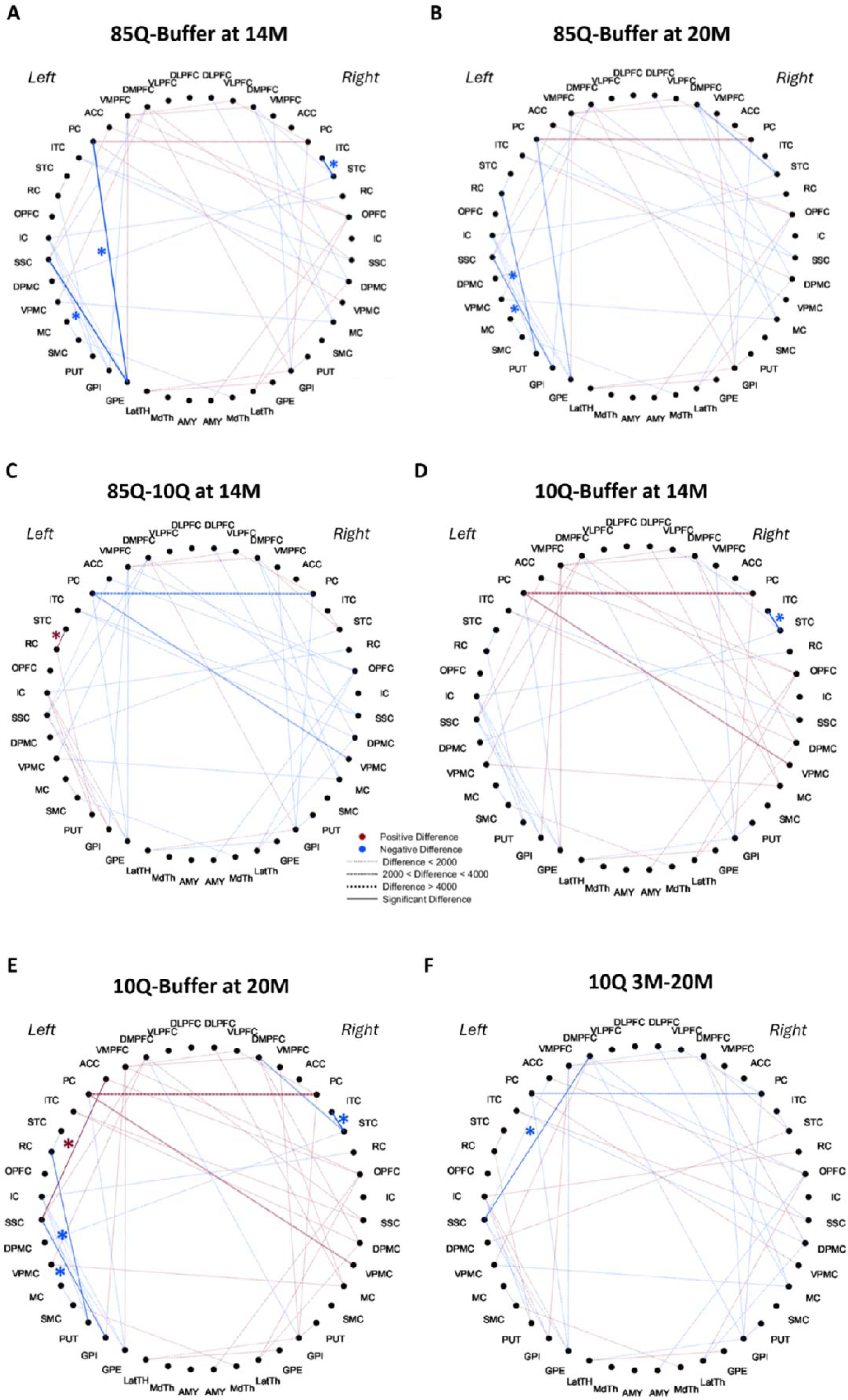
SC differences between Buffer, 85Q and 10Q groups at different timepoints. Circular diagrams illustrating 44 ROIs where a significant interaction effect was found in streamline count (SC) differences between ROIs when comparing the three groups (Buffer, HTT10Qm and 85Q). Dashed lines indicate a significant ANOVA effect while solid lines indicate a significant difference after post-hoc analysis. Blue color indicates a negative difference, red – a positive difference.

**Supplementary Figure 7.**
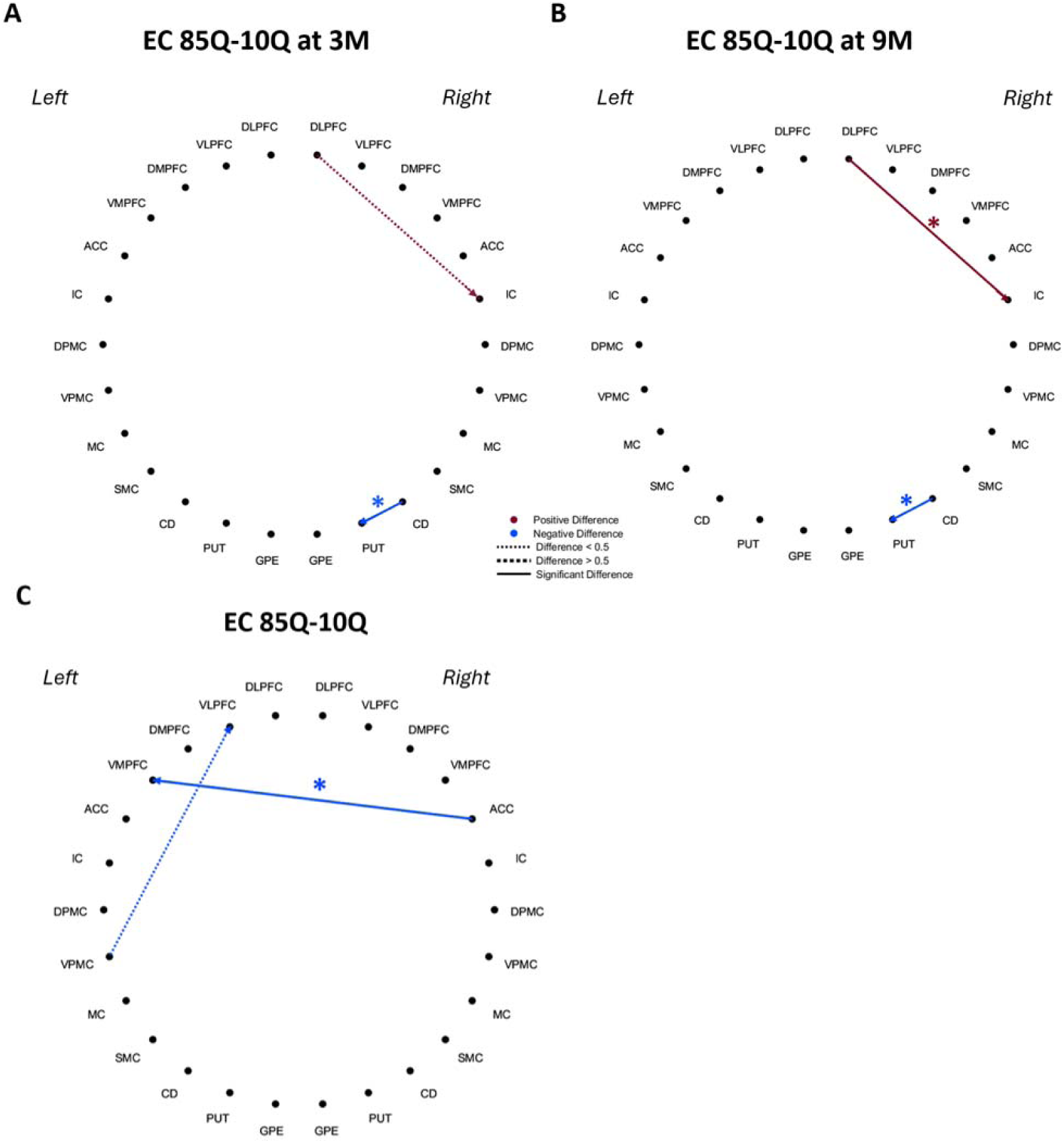
EC comparisons between Buffer, 85Q, and 10Q groups. A, B: A circular diagram illustrating connections (dashed lines) between 26 ROIs where a significant group x time interaction effect was found on EC and those (solid lines with asterisks) that showed significant difference on post-hoc comparisons between 85Q and 10Q at 3 months (A) and at 9 months (B). Blue/red color indicates a lower/higher EC in 85Q as compared to 10Q. C: A circular diagram illustrating difference in EC for connections between 26 ROIs that showed a significant main effect of group on EC (p < 0.05, repeated measures 2-way ANOVA). Blue line indicates a negative difference.

**Supplementary Table 2.** A table containing F-statistic, degrees of freedom and p-values for effective connectivity that showed a significant interaction or group effect followed by a significant difference after post-hoc comparisons.

| Connection Name | Effect Type | Degrees of Freedom | F statistic | P value |
| --- | --- | --- | --- | --- |
| CD to SMC | Interaction (Group x Time) | 1 | 8.12 | 0.0065 |
| PUT to MC | Interaction (Group x Time) | 1 | 6.07 | 0.0161 |
| ACC to VMPFC | Group | 1 | 5.38 | 0.0455 |

